# Acquisition of a type 3 secretion signal in an housekeeping enzyme shaped glycogen metabolism in *Chlamydia*

**DOI:** 10.1101/2020.10.11.335026

**Authors:** Sébastien Triboulet, Maimouna D. N’Gadjaga, Béatrice Niragire, Stephan Köstlbacher, Matthias Horn, Agathe Subtil

## Abstract

The obligate intracellular bacteria *Chlamydia trachomatis* store glycogen in the lumen of the vacuoles in which they grow. Glycogen catabolism generates glucose-1-phosphate (Glc1P), while the bacteria are capable of taking up only glucose-6-phosphate (Glc6P). We tested whether the conversion of Glc1P into Glc6P could be catalyzed by a phosphoglucomutase (PGM) of host or bacterial origin. We found no evidence for the presence of the host enzyme in the vacuole. In *C. trachomatis*, two proteins are potential PGMs. By reconstituting the reaction, and by complementing PGM deficient fibroblasts, we demonstrated that only CT295 displayed robust PGM activity. Furthermore, we showed that glycogen accumulation by a subset of *Chlamydia* species correlated with the presence of a type three secretion (T3S) signal in their PGM. In conclusion, we established that the conversion of Glc1P into Glc6P was accomplished by a bacterial PGM, through the acquisition of a T3S signal in a “housekeeping” gene.

## INTRODUCTION

Many bacteria and parasites develop inside vacuolar compartments within eukaryotic cells. Such enclosed replication niches provide a shelter against extracellular and cytosolic host defense. They can also be further exploited to sequester cytoplasmic components and make them accessible only to the intruder. One striking example of this behavior is the vacuole in which the human adapted pathogen *Chlamydia trachomatis* grows, also called the inclusion (1). Glucose is the main carbon source for these obligate intracellular bacteria (2-5). Over the course of their developmental cycle they hijack a considerable amount of glucose from their host, and accumulate it as a polysaccharide, i.e. glycogen, in the lumen of the inclusion (6, 7). The *C. trachomatis* genome encodes for all the enzymes necessary for a functional glycogenesis and glycogenolysis, a surprising observation given that this pathway is absent in most intracellular bacteria (8). Glycogen starts accumulating in the lumen of the inclusion about 20 hours post infection (hpi), a time of intensive bacterial replication (9). Shortly later, while luminal glycogen continues to accumulate, a fraction of the dividing bacteria, called reticulate bodies (RB), initiate conversion into the non-dividing, infectious, form of the bacterium, the elementary body (EB). Electron microscopy studies showed that, in contrast to RBs, EBs contain glycogen, but in modest quantities compared to the luminal glycogen. While part of the luminal glycogen is imported in bulk from the host cytoplasm, the majority is synthesized *de novo* through the action of bacterial enzymes (9). Bacterial glycogen synthase GlgA, and branching enzyme GlgB, have acquired type 3 secretion (T3S) signals that allow their secretion in the lumen of the inclusion, where they ensure glycogen synthesis out of UDP-glucose imported from the host cytosol (9). T3S of these enzymes is likely shut down once the bacteria have converted into EBs. This simple mechanism might account for the appearance of glycogen in EBs, once GlgA and GlgB can accumulate within the bacteria.

Bacterial enzymes responsible for glycogen catabolism into Glc1P, i.e. the glycogen phosphorylase GlgP, the debranching enzyme GlgX, and the amylomaltase MalQ, have also acquired T3S signals (9). In *C. trachomatis* infection, they are expressed as early as 3-8 hpi, indicating that even if glycogen is only detected 20 hpi, there is a continuous flux of Glc1P generation in the inclusion lumen earlier on (9, 10). However, Glc1P cannot sustain bacterial growth directly because the glucose transporter UhpC can only import Glc6P, not free glucose nor Glc1P (9, 11). Thus, while the rate of expression and distribution of *C. trachomatis* glycogen enzymes and the strategy of glycogen storage in the inclusion lumen by RBs point to a sophisticated pathway for hijacking host glucose, this metabolite would not be able to fuel bacterial metabolism due to a lack of an import mechanism.

To solve this conundrum, we had hypothesized that an enzyme with PGM activity was present in the inclusion lumen (9). In the present work we tested this hypothesis in depth by investigating two non-mutually exclusive possibilities as to its origin: the translocation of a host enzyme into the inclusion, and the secretion of a bacterial enzyme. We provide evidence for the second hypothesis and identify CT295 as *C. trachomatis* PGM. In addition, we show that this feature is characteristic of the glycogen-accumulating *Chlamydiaceae* as opposed to the glycogen-free species.

## RESULTS

### Absence of evidence for the import of host phosphoglucomutase in the inclusion lumen

PGM activity in human cells is carried out by the enzyme PGM1. PGM1 deficiency leads to defects in glycogen storage, and to several disorders resulting from insufficient protein glycosylation (12, 13). To test for the presence of this enzyme in *C. trachomatis* inclusions we expressed PGM1 with either an amino-terminal (Flag-PGM1) or a carboxy-terminal (PGM1-Flag) Flag tag in HeLa cells, which were further infected with *C. trachomatis* L2. The cells were fixed 24 h later and stained with a mouse antibody against the Flag tag and a rabbit antibody against the inclusion protein Cap1. An irrelevant Flag-tagged protein (Flag-CymR) was included as a negative control, and Flag-tagged chlamydial glycogen synthase (Flag-GlgA), as a positive control (9). Flag-PGM1 and PGM1-Flag had a cytosolic distribution, as expected (Fig. 1A). In infected cells, it was often abundant in the vicinity of the inclusion membrane, especially with the carboxy-terminal tag. However, no signal was detected in the inclusion lumen, indicating that PGM1 is not translocated in this compartment. We next tested whether silencing PGM1 expression had any effect on bacterial development. Cells were infected 24 h after siRNA transfection with two different siRNA (si*pgm1*_1 and si*pgm1*_2), and the resulting progeny was measured. The efficacy of each of the siRNA against PGM1 was validated by western blot (Fig. 1B). None of the two siRNA tested had a significant effect on the progeny collected 48 hours post infection (hpi) (Fig. 1C). We concluded from this series of experiments that host PGM1 does not reach the inclusion lumen, at least not in detectable amount, and is not essential for the generation of infectious bacteria. Therefore, it is unlikely that the conversion of Glc1P into Glc6P in the inclusion lumen relies on the host PGM activity by PGM1.

**Fig. 1.**
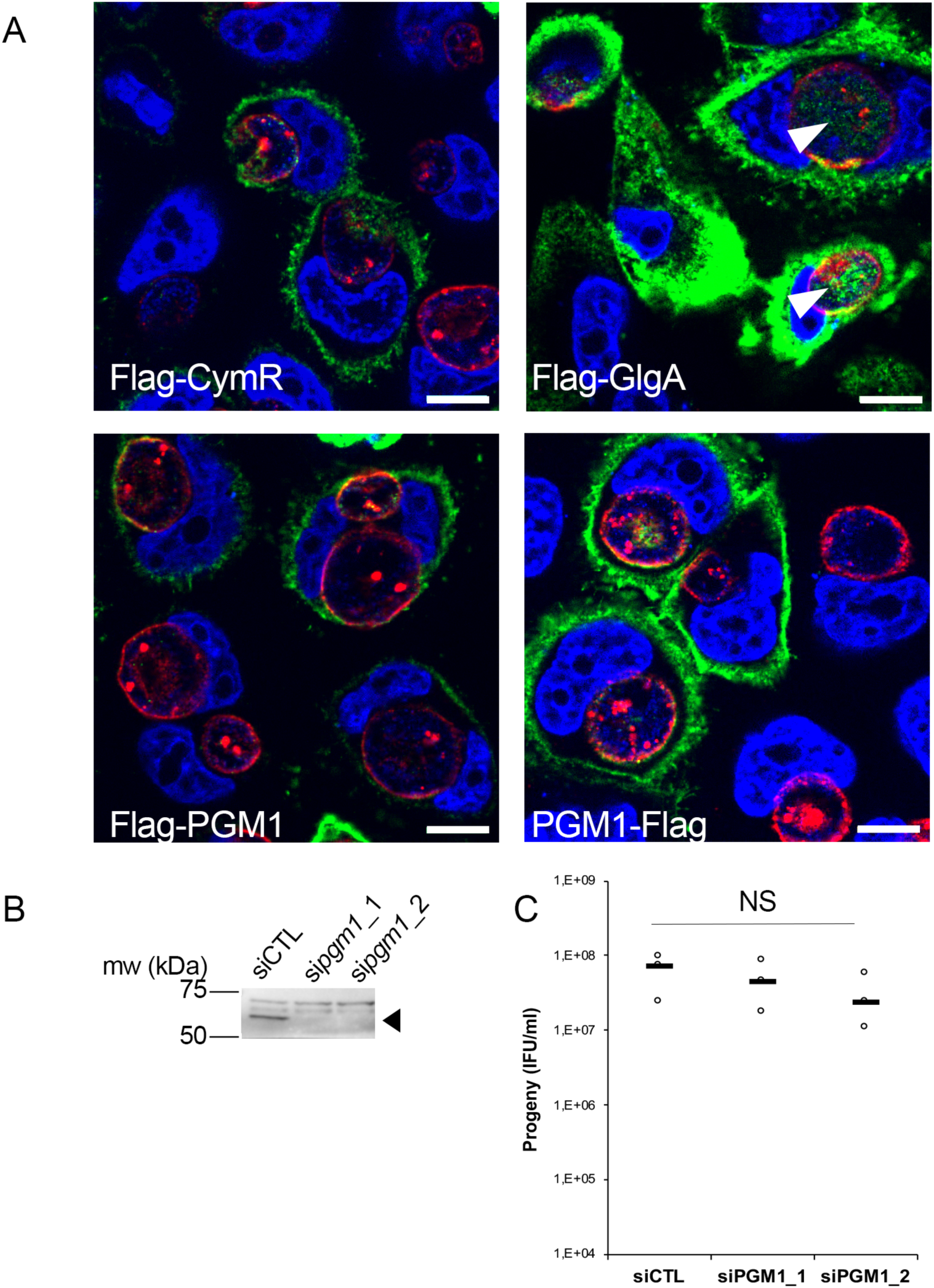
Host PGM1 does not accumulate in the inclusion lumen and is dispensable for *C. trachomatis* development. A – HeLa cells were transfected with the indicated Flag-tagged constructs, infected with *C. trachomatis* L2 and fixed 24 h later. Cells were permeabilized and stained for the inclusion membrane protein Cap1 (red) and Flag (green). DNA was stained with Hoechst (blue). The merge image is shown. Only Flag-GlgA was detected in the inclusion lumen (arrowheads). Bar is 10 μm. B – Two different siRNA were used to silence *pgm1* expression and the effect on PGM1 level was measured by western blot. The arrowhead points to PGM1 (mw=61.5 kDa), higher bands are non specific. C – The effect of *pgm1* silencing on bacterial progeny was measured 48 hpi. Dot plot distribution of the progeny for three independent experiments is shown. Gray bar, median. Statistical significance was calculated using two-way analysis of variance (ANOVA) with Dunnett’s multiple comparison test and was not significant (NS).

### CT295 is the main PGM in *C. trachomatis*, while CT815 displays low activity towards Glc1P

CT295 and CT815 are two *C. trachomatis* proteins that belong to the eggNOG cluster of orthologous groups COG1109 (Fig.2 Figure Suppl.1A, (14)). Members of this large group of bacterial, archaeal and eukaryotic proteins display PGM activity and related functions such as phosphomannomutase or phosphoglucosamine mutase activities. CT295 and CT815 homologs are found among all known chlamydiae. These proteins represent separate monophyletic clades in COG1109 and are related to proteins from diverse bacterial taxa (Fig. S1A). CT295 and CT815 are only distantly related with each other and thus do not originate from a recent gene duplication event (Fig.2 Figure Suppl.1B).

To test if CT295 and/or CT815 displayed PGM activity we expressed them in *Escherichia coli* with an amino-terminal histidine tag. HIS-CT295 was not produced by *E. coli*, and after testing the orthologous protein in different *Chlamydiaceae* we chose the *C. caviae* ortholog, CCA00034, which displayed 59 % amino acid identity with CT295. HIS-CCA00034 and HIS-CT815 were purified from *E. coli* extracts (Fig.2 Figure Suppl.2A). As a positive control we also produced the *E. coli* phosphomannomutase CpsG, because our initial attempts to produce the *E. coli* PGM failed, and CpsG turned out to display good PGM activity *in vitro*. To test the ability of these recombinant proteins to convert Glc1P into Glc6P we first incubated them with Glc1P in the presence of glucose 1,6-diPhosphate (Glc1,6diP), a cofactor for the PGM reaction in *E. coli* (15). The samples were incubated for 20 min at 37 °C, before stopping the reaction by boiling for 5 min. Sugars present in the samples were analyzed by high performance anion exchange chromatography with pulsed amperometric detection (HPAEC-PAD) (Fig.2 Figure Suppl.2B). Incubation of Glc1P in the presence of HIS-CCA00034 led to its full conversion into Glc6P, while only low levels of Glc6P were detected in the presence of HIS-CT815 (Fig. 2A). Interestingly we noted that while we confirmed that HIS-CpsG required the presence of Glc1,6diP to convert Glc1P into Glc6P, HIS-CCA00034 did not, indicating that the chlamydial enzyme evolved to bypass co-factor requirement (Fig.2 Figure Suppl.2C). Also, HIS-CCA00034 favored the conversion of α-Glc6P into β-Glc6P, a property that we did not observed in CT815 nor CpsG (Fig. 2 and Fig. Suppl.2C-D).

**Fig. 2.**
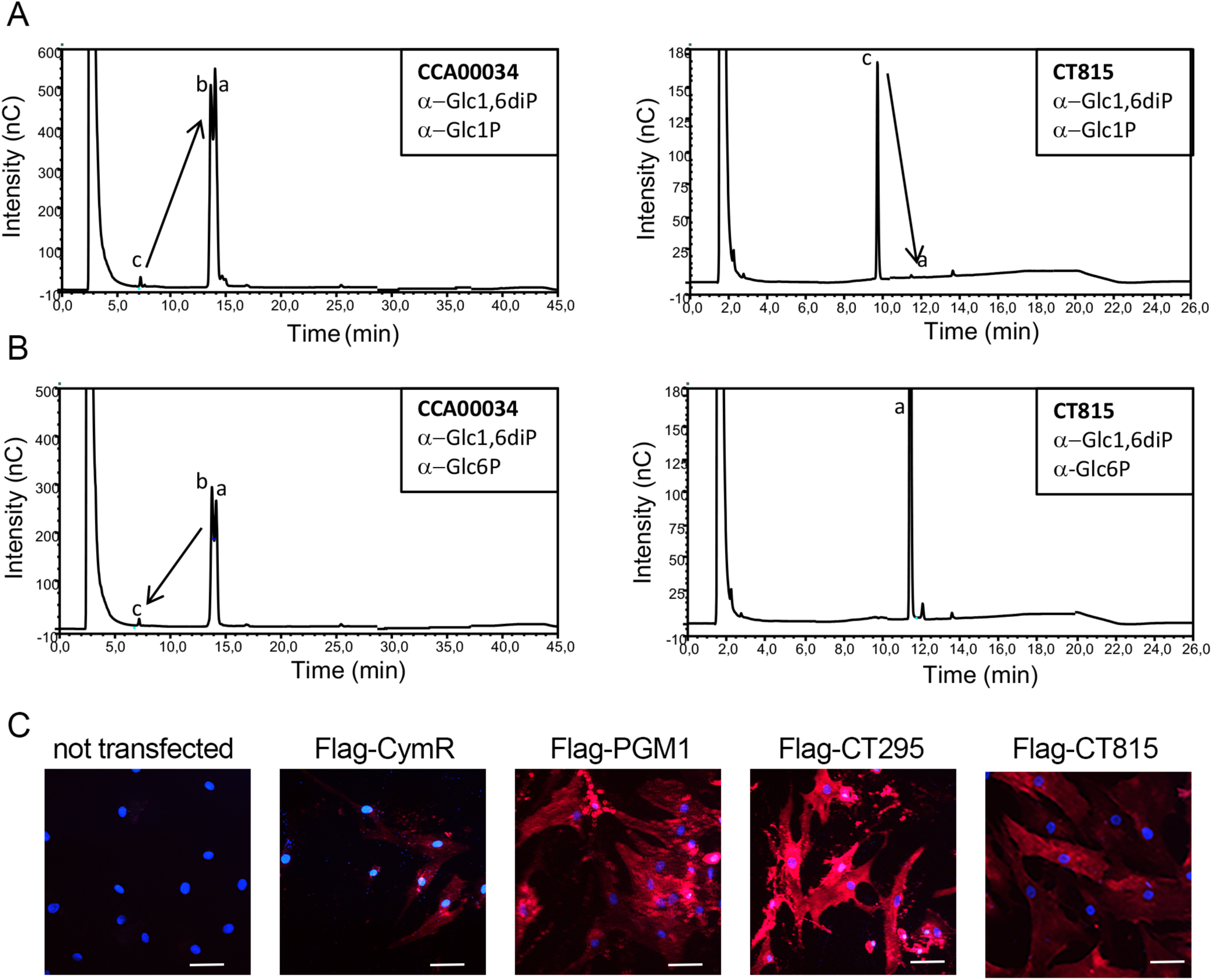
CT295 is a phosphoglucomutase. A – The indicated purified proteins (bold letters) were incubated with Glc1P in the presence of the co-factor Glc1,6diP. The substrates and reaction products were identified by HPAEC. Incubation of Glc1P (peak c) with CCA00034 resulted in full conversion into Glc6P (peaks a and b), while Glc6P was hardly detectable when CT815 was tested. Note that the samples were run on two different columns, resulting in different elution profiles. B – The reverse reaction was tested. A very low conversion of Glc6P into Glc1P was observed for CCA00034 and was totally absent for CT815. Of note, CCA00034 favors the conversion of α-Glc6P (peak a) into β-Glc6P (peak b) while CT815 does not. C – PGM1 deficient fibroblasts were transfected or not with the indicated constructs. One day later, ICAM1 expression at the cell surface was probed with Cy5-coupled anti-ICAM1 antibody (red). Nuclei were stained with Hoechst (blue). Bar is 50 μm.

When we monitored the reverse reaction, supplying Glc6P and following the appearance of Glc1P, very low amounts of Glc1P were detected after incubation with HIS-CCA00034 or HIS-CpsG, and none with HIS-CT815 (Fig. 2B and Fig. Suppl.2D). This indicates that, *in vitro*, the conversion of Glc1P into Glc6P is more favorable than the reverse reaction. To confirm that CT295 was nonetheless capable of catalyzing the reverse reaction, we made use of human fibroblasts deficient for PGM1. One of the phenotypes associated with PGM1 deficiency is a defect in protein glycosylation, due to the inability to convert Glc6P into Glc1P to feed UDP-Glc synthesis and thereby protein glycosylation (13). As a consequence, several glycosylated proteins like the cell-surface glycoprotein intercellular adhesion molecule 1 (ICAM-1) are hardly expressed at the cell surface in PGM1 deficient cells (13). ICAM-1 expression can thus be used as a read-out for PGM activity. As expected, ICAM-1 was hardly detected at the surface of PGM1 deficient fibroblasts. Transfection of an irrelevant protein, Flag-CymR, had a marginal effect on ICAM-1 expression. In contrast, as expected, transfection of the wild-type human PGM1 restored ICAM1 expression in the PGM1 deficient fibroblasts. Transfection of CT295 also restored ICAM-1 expression, confirming its ability to catalyze the conversion of Glc6P into Glc1P. Transfection of CT815 also resulted in increased ICAM-1 expression, but to a lesser extent, supporting the hypothesis that while CT815 displays some phosphoglucomutase activity, it is likely that it uses a different sugar than glucose as a favorite substrate (Fig. 2C).

Altogether, these data demonstrate that CT295, and its ortholog in *C. caviae*, are bona fide phosphoglucomutases.

### CT295 is a T3S substrate

If CT295 is the PGM that converts Glc1P into Glc6P in the inclusion lumen it needs to be secreted out of the bacteria. We have shown that *C. trachomatis* glycogen metabolism enzymes were secreted by a T3S mechanism (9). To explore if this was also true for CT295, we tested for the presence of a functional T3S signal within its amino-terminus using a heterologous secretion assay in *Shigella flexneri* as described previously (16). Briefly, we fused these amino-terminal sequences to the reporter protein calmodulin-dependent adenylate cyclase (Cya). The chimeric proteins were then expressed in *S. flexneri ipaB* (constitutive T3S) or *mxiD* (deficient in T3S) null strains. When we analyzed the culture supernatant versus the bacterial pellet, we found the Cya reporter in the supernatant of *ipaB* cultures (Fig. 3A). The same expression pattern was observed for an endogenous *Shigella* T3S substrate, IpaD. Conversely, we found the cAMP receptor protein (CRP), a non-secreted protein, exclusively in the bacterial pellet. This control shows that Cya detection in the supernatant was not due to bacterial lysis. Finally, the Cya reporter was not recovered in the supernatants of the *mxiD* strain, demonstrating that its secretion was dependent on the T3S system.

**Fig. 3.**
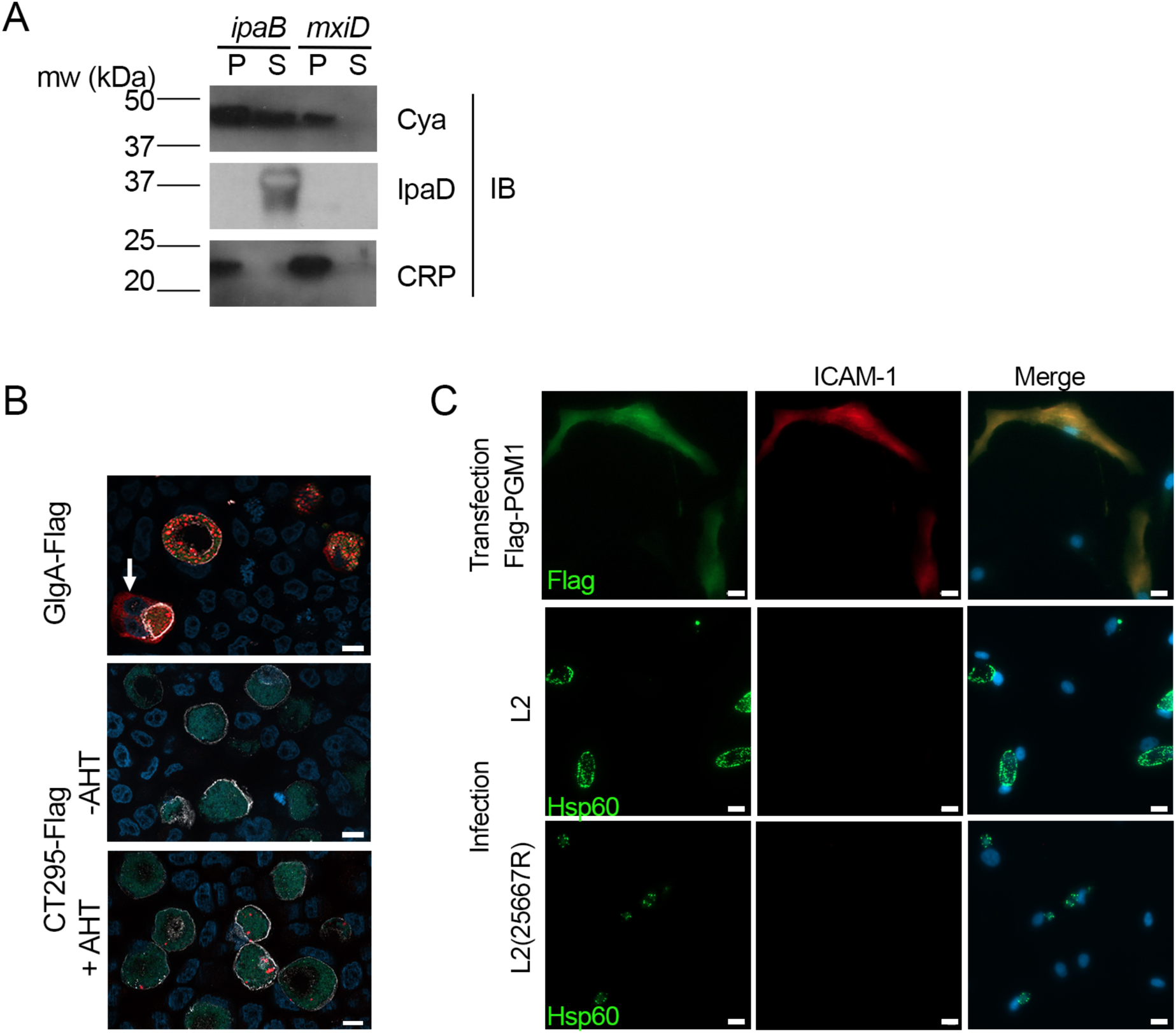
CT295 is secreted in the inclusion lumen. A – The first 22 amino acids of CT295 were fused to the Cya reporter protein and expressed in a S. *flexneri ipaB* (constitutive T3S) or *mxiD* (defective T3S) strain. Exponential phase cultures expressing the reporter fusion protein were fractionated into supernatants (S) and pellets (P). Samples were resolved by SDS-PAGE, transferred to a PVDF membrane, and probed with anti-Cya (to detect the fusion protein), anti-IpaD (*Shigella* secreted protein), or anti-CRP (*Shigella* non-secreted protein) antibodies. B – Cells were infected with *C. trachomatis* L2 strain stably expressing GFP together with either GlgA-Flag (endogenous promoter), or CT295-Flag (Tet-inducible promoter), and incubated in the absence of presence of 5 nM anhydrotetracycline (AHT) for 48 h before fixation, permeabilization and staining. Bacteria are visible in the green channel, the inclusion membrane is labeled with antibodies against Cap1 followed with Cy3-coupled anti-rabbit antibodies (white), secondaries antibodies for the Flag-tagged were Alexa647-coupled anti-mouse antibodies (red), the DNA was stained with Hoechst (blue). The white arrow points to GlgA-Flag detected in the cytoplasm of one infected cell. Bar is 10 μm. C – PGM1 deficient fibroblasts were transfected with Flag-PGM1 or infected with the indicated *C. trachomatis* L2 strain. Forty-eight hours later the cells were fixed, permeabilized and stained with the indicated antibodies. Nuclei were stained with Hoechst (blue). Bar is 20 μm.

### Evidence for the direct secretion of CT295 in the inclusion lumen

*C. trachomatis* GlgA is detected both in the inclusion lumen and in the cytoplasm (17). When expressed ectopically by the host in the cytoplasm, this bacterial enzyme was able to translocate into the inclusion lumen (9). These observations suggested that some T3S substrates might be first translocated in the cytoplasm, before being transferred into the inclusion lumen. To test whether this was also the case for CT295, we first transformed *C. trachomatis* with an expression plasmid constitutively expressing the green fluorescent protein (GFP), as well as CT295 with a carboxy-terminal Flag tag under the control of an inducible promoter, as an initial attempt using the endogenous promoter showed no protein expression. Bacteria expressing GlgA-Flag under the control of its endogenous promoter were used for comparison. The cells were fixed 48 hpi, permeabilized, and stained with a mouse antibody against the Flag tag and a rabbit antibody against the inclusion membrane protein Cap1. As expected, a cytoplasmic staining was observed for GlgA-Flag, although only in ∼ 10% of the infected cells (n=150 infected cells from 2 independent experiments). A faint Flag staining was detected in the inclusions upon induction of cells infected with bacteria expressing CT295-Flag (Fig. 3B). No signal was detected in the host cell cytoplasm, indicating that the protein is not translocated there. However, the expression level of CT295-Flag was much lower than that of GlgA-Flag, whose cytoplasmic location is only visible in ∼ 10% of the cells. Thus, this assay might not be sensitive enough to detect CT295 in the cytoplasm.

As an alternative approach to test for CT295 presence in the host cytoplasm we tested whether infection could restore PGM activity in the cytoplasm of PGM1 deficient fibroblasts. Indeed, if CT295 reached the cytoplasm, it might lead to at least a partial recovery of ICAM-1 glycosylation. Cells transfected with Flag-PGM1 served as a positive control. We observed that ICAM-1 expression, which was undetectable at the surface of PGM1 deficient fibroblast, remained absent from the surface of cells infected for 40 h with *C. trachomatis* (Fig. 3C). One pitfall of this experiment is that *C. trachomatis* L2 takes up UDP-glucose from the host cytoplasm to accumulate glycogen in the inclusion lumen (9). Therefore, the presence of a bacterial PGM in the cytoplasm might not be sufficient to restore ICAM-1 glycosylation. We thus also used the plasmid-less L2(25667R), which is less likely to deplete the host UDP-glucose stores, as it does not accumulate glycogen in the inclusion (18). Infection with L2(25667R) did not restore ICAM-1expression at the cell surface either. These experiments indicate that CT295 is translocated directly into the inclusion lumen, without an intermediate cytoplasmic step (Fig. 3C).

### Presence of a T3S in CT295 orthologs among *Chlamydiaceae* correlates with their capacity to accumulate glycogen in the inclusion lumen

We next asked whether the secretion of a PGM was a conserved feature among *Chlamydiaceae*. Previous studies have reported glycogen accumulation in *C. suis* and *C. muridarum* inclusions, but the polysaccharide was not observed in the inclusions of more phylogenetically distant *Chlamydia* species such as *C. caviae* or *C. pneumoniae*. We verified this feature by performing periodic acid-Schiff staining (PAS) on cells infected with these different species. Inclusion staining was observed only on cells infected with *C. trachomatis, C. suis* and *C. muridarum* (Fig. 4A). Two amino-terminal amino-acids (W23L24) are conserved in the orthologs of CT295 in all chlamydiae, and the preceding amino acids show low levels of conservation (Fig. 4B). To test whether these different peptides also function as T3S signals in other species than *C. trachomatis* we constructed chimeras between the amino-terminal 22 amino-acids of the different CT295 orthologs and the Cya reporter gene, and tested their secretion when expressed in the *S. flexneri ipaB* strain. The chimera made out of the amino-terminal segment of *C. suis* and *C. muriarum* PGMs were recovered in the supernatant, but not those originating from *C. caviae* or *C. pneumoniae*. Secretion was dependent on the T3S system since the chimera were not recovered in the supernatant when expressed by the *S. flexneri mxiD* strain (Fig. 4C). Thus, the accumulation of glycogen in the inclusion of *Chlamydiaceae* correlates with their ability to secrete a phosphoglucomutase.

**Fig. 4.**
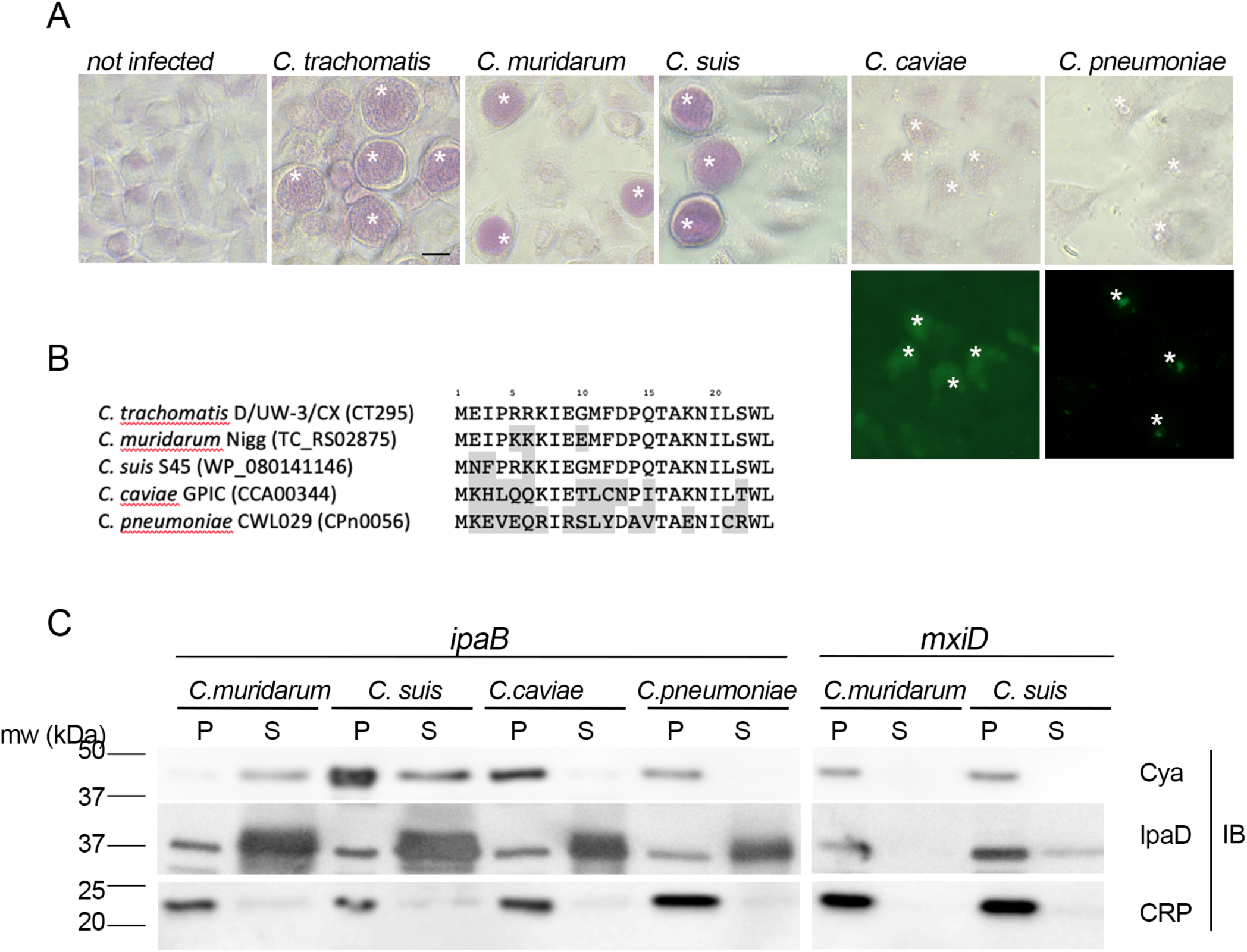
Secretion of a PGM activity correlates with glycogen accumulation among *Chlamydiaceae*. A – Top panels: PAS staining of cells infected with the indicated strains. Bottom panels: *C. caviae* and *C. pneumoniae* infected cells were further stained with anti-mGroEL antibody followed with Alexa488-conjugated secondary antibodies to detect the inclusions (asterisks). Bar is 20 μm. B – Alignement of the first 24 amino acids of CT295 and its orthologs in the indicated *Chlamydiaceae* species. Amino acids that differ from the *C. trachomatis* sequence are shadowed. C – The first 22 amino acids of the indicated CT295 orthologs were fused to the Cya reporter protein and expressed in a S. *flexneri ipaB* (left) or *mxiD* (right) strains. Secretion was analyzed like in Fig. 3A.

## DISCUSSION

To be of metabolic use for the bacteria the glycogen stored in the inclusion lumen needs to be converted into glucose moieties. Glycogen degradation in the inclusion generates Glc1P, and the single glucose transporter of *C. trachomatis*, UhpC, displays transport activity exclusively towards Glc6P. We show here that this conundrum is solved by the secretion of a bacterial PGM, CT295, by the T3S system. Host PGM1 was not translocated into the inclusion lumen, at least not at levels detectable by immunofluorescence. Silencing *pgm1* did not affect the generation of infectious bacteria. Thus, while we cannot exclude that imported host PGM1 also contributes to Glc1P to Glc6P conversion in the inclusion lumen, our data suggest that such activity would only be marginal compared to that of secreted CT295.

*In vitro* and *in cellulo* assays demonstrated that CT295 and its ortholog in *C. caviae* displayed *bona fide* PGM activity. Interestingly, unlike what had been observed in *E. coli* (15), the chlamydial activity did not require the co-factor glucose 1,6-diPhosphate (Glc1,6diP). This may represent an adaptation to the inclusion environment, where this metabolite might be absent. The second *C. trachomatis* enzyme with predicted PGM activity, CT815, diverged from CT295 early in the evolution of the phylum, possibly even before its emergence. It displayed a poor ability to interconvert Glc1P and Glc6P *in vitro* and *in vivo*, indicating that its preferred substrate is different, possibly phosphoglucosamine, as shown for at least one member or the same clade (19).

Transcriptomic studies detected transcription of *ct295* 8 hpi, a timing that coincides with the start of expression of the Glc6P transporter *uhpC* (10). Interestingly, the protein CT295 was hardly detected by proteomics in the replicative form of the bacteria, the reticulate bodies (RBs), while it was rather abundant in the infectious form, the elementary bodies (EBs) that accumulate at the end of the developmental cycle (20). This difference in abundance in the two bacterial forms is consistent with the observation that glycogen is detectable in EBs, not in RBs (9). It might be explained by the fact that most CT295 synthesized by RBs is secreted to convert Glc1P to Glc6P in the inclusion lumen, and thus absent from purified RBs. Like other T3S substrates, secretion of CT295 probably stops during conversion to the EB stage, and the enzymatic activity is then mostly exerted inside the bacteria. Here, it is likely used for the conversion of Glc6P into Glc1P, which serves as a substrate for the ADP-Glc pyrophosphorylase GlgC to initiate glycogen synthesis in the bacteria.

We and others have shown that several T3S substrates are detected in the inclusion lumen (9, 21, 22). This observation was at first surprising considering that most T3S machineries have been described to deliver proteins across a eukaryotic membrane, through the translocon pore. Here, we considered the possibility that CT295 was first translocated into the cytoplasm before being transported back inside the inclusion lumen. However, we found no evidence of an increase in PGM activity in the host cytoplasm upon infection. It is therefore likely that CT295 is secreted directly into the lumen of the inclusion. Of note, several T3S substrates have been detected on the surface of different Gram-negative bacteria before being translocated into a mammalian cell (23, 24). These observations support the hypothesis of direct secretion of T3S substrates into the inclusion lumen rather than via the T3S translocon pore into the host cell. Future work will be needed to understand how these substrates are sorted from those that are translocated into the cytoplasm.

Finally, we made the intriguing observation that chlamydial PGMs possess functional T3S signals only in species that accumulate glycogen in the inclusion lumen. These species are phylogenetically apart from other *Chlamydiaceae*, and differ by several other features (25). We lack sufficient information on the translocation, in the inclusion lumen or in the host cytoplasm, of glycogen metabolism enzymes of *Chlamydiaceae* other than *C. trachomatis* to draw solid hypotheses on the evolutionary path that led to this situation, we can however try to elaborate on this observation. We have shown that T3S signals were present in glycogen metabolism enzymes of several environmental chlamydiae, indicating that the ability to take control of glycogen storage capacity of the host was acquired early upon evolution of the phylum (26, 27). One possibility is that the acquisition of T3S signal in the PGM of the *Chlamydia* lineage (as opposed to species of the formerly called *Chlamydophila* lineage, e.g. *C. pneumoniae* and *C. Caviae*) opened the way to a positive selection of the capacity to store glycogen in the inclusion. Indeed, once Glc1P became convertible to Glc6P, it might have become advantageous to store a lot of glycogen, possibly to compete out the rest of the microbiota, or to weaken host cell defense, as previously discussed (9). Of note, expression of the glycogen synthase GlgA is the limiting factor for glycogen accumulation in the inclusion. In *C. trachomatis*, this gene is under transcriptional control by the *C. trachomatis* plasmid (18). Acquisition of this transcriptional control was likely also an important step along the path towards the glycogen storage strategy.

One question left open is whether a more direct mechanism for Glc6P supply in the inclusion lumen of *C. trachomatis* also takes place? An alternative mechanism must exist at least in the *Chlamydiaceae* of the former *Chlamydophila* lineage, considering the absence of glycogen and phosphoglucomutase in their inclusions, and the fact that UhpC transports only Glc6P also in *C. pneumoniae* (11). The situation is different in environmental chlamydiae because, unlike *Chlamydiaceae*, environmental chlamydiae express a hexokinase activity (28-30), and are likely able to import glucose, as demonstrated in the case of *Protochlamydia amoebophila* (31).

In conclusion, we have shown that several “housekeeping” chlamydial proteins are secreted outside the bacteria. This possibility has seldom been investigated in other bacteria, but was also observed for glyceraldehyde-3-phosphate dehydrogenase in enteropathogenic *E. coli* (32). In that case, the broad range type 3 secretion chaperone CesT was involved, and whether such a chaperone is required for the secretion of chlamydial enzymes remains to be investigated. In any case, the secretion of bacterial enzymes opens the door to a direct modification of the metabolism of the host cell to the benefit of the bacteria, and might therefore occur more frequently than currently known.

## MATERIAL AND METHODS

### Cells and bacteria

HeLa cells (ATCC) were cultured in Dulbecco’s modified Eagle’s medium (DMEM) with Glutamax (Gibco™ #31966), supplemented with 10 % (v/v) fetal bovine serum (FBS). Fibroblast CDG_0072 deficient in PGM1, a kind gift from Pr. Hudson Freeze (La Jolla, CA), were grown in DMEM (Gibco™ # A1443001) supplemented with 1 g/L glucose, 584 mg/mL L-glutamine, 110 mg/mL sodium pyruvate, and 10% FBS. Cells were routinely checked for absence of mycoplasma contamination by PCR. *C. trachomatis* LGV serovar L2 strain 434/Bu (ATCC), the plasmid-less strain LGV L2 25667R (33) and GFP-expressing L2 (L2^IncD^GFP) (34) were purified on density gradients as previously described (35). Other *Chlamydia* strains used were *C. suis* SWA107, a kind gift from Nicole Borel (Zürich, Switzerand), *C. pneumoniae* CWL029, *C. muridarum* MoPn and *C. caviae* GPIC. The *Shigella flexneri ipaB* and *mxiD* strains are derivates of M90T, the virulent wild-type strain, in which the respective genes (*ipaB* and *mxiD*) have been inactivated (36). The *Escherichia coli* strain *DH5*α was used for cloning purposes and plasmid amplification. Both *S. flexneri* and *E. coli* strains were grown in Luria-Bertani (LB) medium.

### Phylogenetic analysis of CT295 and CT815

Protein sequences of CT295 and CT815 were analyzed with eggNOG v4.5.1 (14). They are both members of COG1109 at the last universal common ancestor (LUCA) taxonomic level (including all domains of life). At the bacteria-specific eggNOG clustering, CT295 and CT815 belong to two distinct EggNOGs, ENOG4107QSU and ENOG4107QJF. To identify closely related sequences we used eggNOG-mapper v1.0.1 (37) with the bacteria optimized database using the “--database bact” option and default settings on a broad selection of published chlamydial genomes. We downloaded all COG1109 sequences (n=4,829; http://eggnogapi45.embl.de/nog_data/text/fasta/COG1109), added all chlamydial ENOG4107QSU (n=104) and ENOG4107QJF (n=69) sequences and aligned them using the COG1109 hidden markov model (http://eggnogapi45.embl.de/nog_data/file/hmm/COG1109). To reduce redundancy we clustered the alignment with the cd-hit program (38) at 80% sequence identity over at least 90% of the shorter sequence (“-aS 0.9 -c 0.8”). We performed de novo multiple sequence alignment of the clustered sequences (n=2,869) with MAFFT 7.222 (39) using the “--localpair” and “--maxiterate 1000” parameter and trimmed the alignment with trimAl “-gappyout” (40). Only sequences with a gap-rate < 50% were retained (n=2,829). We performed model testing under the empirical LG model (41), and with the empirical mixture models C10 to C60 (best model: C60+LG+G+F) and maximum likelihood tree reconstruction with IQ-TREE 1.6.2 (42). We inferred support values from 1,000 ultrafast bootstrap replicates (43) with the “-bnni” option for bootstrap tree optimization and from 1,000 replicates of the SH-like approximate likelihood ratio test (44).We next extracted the sister clade of CT295 (n=93) and used it as an outgroup for a more detailed analysis of CT295 related proteins. Sequence alignment, trimming, and initial phylogenetic reconstruction (best model: C60+LG+G+F) was performed as described above. We then used the resulting best tree as a guide tree for the posterior mean site frequency (PMSF) model (45) for improved site heterogeneity modeling under C60+LG+G+F and inferred 100 non-parametric bootstraps. Phylogenetic trees were visualized and edited using the Interactive Tree Of Life v4 (46).

### Cloning

Primer sequences and cloning methods are described in Supplementary file 1, all constructs were verified by sequencing. The gene *ct815* was cloned as follows: the gene was amplified with primers 1 and 4, and with primers 2 and 3. PCR products were mixed, heated to 65 °C for 10 minutes, and annealed at 37 °C for 30 minutes. The mix was then cloned into the pET28a previously digested by NcoI and XhoI restriction enzymes. Chimeras to test for the presence of T3S signal were constructed like in (16), using the first 22 codons of the gene of interest. Flag-GlgA is described in (9) and the negative control, CymR, that corresponds to a bacterial repressor for gene expression from *Pseudomonas putida* (47) was constructed through Gateway cloning like Flag-PGM1 and PGM1-Flag. The *ctl0547* gene (CT295 in the *C. trachomatis* L2/434/Bu strain) followed by a nucleotide sequence coding for a flag tag immediately upstream of the stop codon was cloned using NotI and SalI restriction sites into the pBOMB4-Tet-mCherry plasmid after removing the mCherry insert (48). For GlgA, cloning was performed in pBOMB4 using the endogenous promoter of the *ctl0167* gene. The strain LGV L2 25667R was stably transformed with these plasmid as described (49).

### Production and purification of recombinant proteins

*E. coli* BL21 (DE3) was used for the production of recombinant CCA00034, CT815, and CpsG harboring a polyhistidine tag at the C-terminal part of the protein. Bacteria were grown at 37 °C to an optical density at 600 nm of 0.9 in 2 liters of LB with 30 µg/mL of kanamycin under shaking. 0.5 mM of isopropyl-β-D-thiogalactopyranoside (IPTG) was added and incubation was continued for 18 h at 16 °C. Bacteria were harvested by centrifugation, and proteins were purified from clarified lysate by affinity chromatography on Ni^2+^-nitrilotriaceta-agarose (Qiagen) in 20 mM Tris-HCl 500 mM NaCl pH 8.0 buffer and by size exclusion chromatography on Superdex 200 HL 16/600 column (GE Healthcare) in 20 mM Tris-HCl 500 mM NaCl pH 7.0 buffer. All purified proteins were stored at 4 °C.

### Phosphoglucomutase activity assay

HIS-tagged CCA00034, CT815 or CpsG (0,1 mg/mL) were incubated for 20 minutes at 37 °C with 0.2 mg/mL of glucose-1,6-diphosphate and either 0.66 mg/ml glucose-1-phosphate or 0.33 mg/ml glucose-6-phosphate in 10 mM Tris-HCl buffer (pH 8.0) and then boiled for 5 minutes to stop the reaction. Reaction products were analyzed by High Performance Anion Exchange Chromatography with Pulse Amperometric Detection (Dionex, model ICS3000). The samples were loaded on a CarboPAC-PA100 column (Dionex) pre-equilibrated for 20 minutes with 98% of 50 mM NaOH (eluant A) and 2% of 50 mM NaOH 500 mM NaOAc (eluant B). After injection a gradient run (flow rate of 0.350 mL.min^-1^) was performed as follows: 0-2 min 98% A + 2% B – 88.8% A + 11.2% B, 2-20 min 88.8% A + 11.2% B – 65% A + 35% B, 20-35 min 65% A + 35% B – 40% A + 60% B, 35-37 min 40% A + 60% B – 100% B. The data displayed are representative of at least 2 independent enzymatic assays.

### Transfections, immunofluorescence and PAS staining

Cells were transfected with the indicated plasmids using the JetPrime kit (Polyplus Transfection) according the manufacturer recommendations. Twenty-four hours post transfection, cells were fixed with 2% (w/v) paraformaldehyde (PFA) 2% (w/v) sucrose in PBS for 30 minutes at room temperature, then washed with PBS twice. For the panel shown in Fig.1, a 4% PFA 4 % sucrose solution was used for fixation. Cells were blocked with 10 mg/mL bovine serum albumin (BSA) for 30 minutes before being stained with antibodies anti-ICAM-1 coupled to APC (Invitrogen 17.0549.42) at dilution 1/1000. Cells were then washed twice with PBS, permeabilized with 0.5% saponin in PBS and DNA was stained with Hoechst 33342 (Molecular Probes). For the panel shown in Fig. 3D the cells were permeabilized after ICAM-1-APC staining to probe for Flag (rabbit, Sigma F7425) or Hsp60 (rabbit, homemade) followed with A488-conjugated secondary antibodies. For the panel shown in Fig.1 and 3C, the cells were permeabilized in 0.5% saponin in PBS and stained with mouse anti-Flag M2 antibody (Sigma) and homemade rabbit anti-Cap1 antibody, followed with species-specific secondary antibodies. Images were acquired on an Axio observer Z1 microscope equipped with an ApoTome module (Zeiss, Germany) and a 63× Apochromat lens. Images were taken with an ORCA-flash4.OLT camera (Hamamatsu, Japan) using the software Zen. All images shown are representative of observations made on at least 3 biological replicates, in which more than 50 cells were observed each time.

For periodic-acid-Schiff (PAS) stain cells were fixed in 4 % PFA 4% sucrose in PBS for 30 min at room temperature and staining was performed as described (50). Briefly, cells were incubated in 1 % periodic acid (Sigma) for 5 min. Thereafter coverslips were put in tap water for 1 min, quickly rinsed in mQ-H_2_0 and then applied to Schiff reagent (Sigma) for 15 min at room temperature. Afterwards the coverslips were rinsed again in mQ-H_2_0, incubated in tap water for 10 min followed by an incubation step in PBS for 5 min. Periodic acid oxidizes the vicinal diols in sugars such as glycogen to aldehydes, which now in turn react with the Schiff reagent to give a purple colour. Coverslips infected with *C. caviae* and *C. pneumoniae* were further stained with home-made rabbit antibody against *C. muridarum* GroEL, that cross-reacts with the orthologous proteins in these species, followed with Alexa488-conjugated secondary antibodies. Images were acquired using transmission light or green fluorescence excitation on an Axio Vert.A1 microscope (Zeiss, Germany) and an LD A-Plan 40x lense. Images were taken with an AxioCam ICc 1 camera using the software Zen. All images shown are representative of observations made on at least 2 biological replicates, in which more than 50 inclusions were observed each time.

### Progeny assays

For siRNA transfection 50 000 cells were plated in a 24-well plate and immediately mixed with Lipofectamine RNAiMAX (Invitrogen) following the manufacturer’s recommendation, using 10 nM of siRNA (see Supplementary file 1 for sequences). Cells treated with siRNA for 24 h (J1) were infected with L2^IncD^GFP bacteria at a MOI = 0.2. Transfection of siRNAs was repeated on J2. On J3 (48 hpi), cells were detached, lysed using glass beads and the supernatant was used to infect new untreated cells plated the day before (100 000 cells/well in a 24-well plate), in serial dilution. The next day, 3 wells per condition with an infection lower than 30 % (checked by microscopy) were detached, fixed and analyzed by flow cytometry to determine the bacterial titer as described (34).

### *Shigella* heterologous secretion assays and immunoblots

Chimera were constructed as described previously (16) with the primers and templates displayed in Supplementary file 1. The sequence of all chimera was verified by sequencing. Analysis of secreted proteins was performed as described previously (16), except that the *ipaB* or *mxiD Shigella flexneri* were co-transformed with the pMM100 plasmid that codes for the *lacI* repressor (selection with tetracyclin) (51) and with the indicated Cya chimera (selection with ampicillin). This was done to avoid the very strong expression of some of the chimeras in the absence of *lacI* repressor. Briefly, 1 ml of a 30 °C overnight culture of *S. flexneri ipaB* or *mxiD* transformed with different Cya chimeras was inoculated in 15 ml of LB broth and incubated at 37 °C for 1 h. Expression of the Cya chimeras was induced by adding 10 µM IPTG for all chimera except PGMtracho/Cya which was induced at 100 µM IPTG. Bacteria were harvested by centrifugation 3 h later and the supernatant was filtered through a Millipore filter (0.2 μm). To precipitate the proteins 1/10 (v/v) of trichloroacetic acid was added to the supernatants. The pellet fraction and the supernatant fraction (concentrated 40-fold compared to the pellet fraction) were resuspended in sample buffer for SDS-PAGE, transferred to Immobilon-P (PVDF) membranes and immunoblotted with the proper primary antibodies diluted in 1X PBS containing 5% milk and 0.01% Tween-20. Primary antibodies used were mouse anti-cya, rabbit anti-CRP and rabbit anti-IpaD antibodies generously given by Drs N. Guiso, A. Ullmann and C. Parsot, respectively (Institut Pasteur, Paris). Goat anti-mouse and anti-rabbit IgG-HRP (GE Healthcare) were used at 1:10000 dilution. Blots were developed using the Western Lightning Chemiluminescence Reagent (GE Healthcare). The data displayed are representative of at least three independent experiments.

## ACKNOWLEDGEMENTS

We thank Dr. Hudson Freeze (Sanford-Burnham Medical Research Institute, La Jolla, USA) for the generous gift of PGM1-deficient fibroblasts, Dr. Hanna Marti (University of Zürich) for the *C. suis* strain and Dr. Lena Gehre for the GlgA-Flag expressing bacteria. This work was supported by the Agence Nationale pour la Recherche (ANR-14-CE11-0024-02 “Expendo”), the Institut Pasteur and the Centre National de la Recherche Scientifique. D.N’G. received financial support from the Fondation pour la recherche médicale (FDT202012010504). MH received financial support from the Austrian Science Fund FWF (DOC 69-B).

---

**Figure 2: Figure supplement 1.**
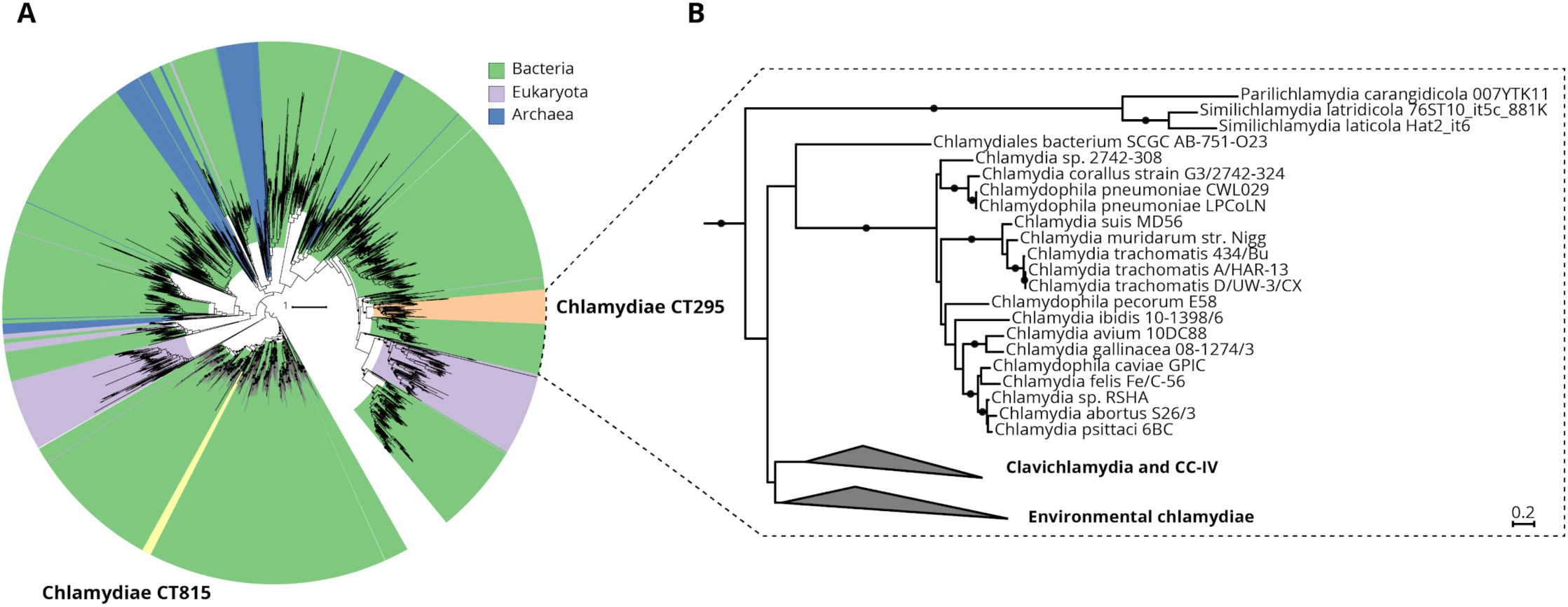
The chlamydial candidate PGMs CT295 and CT815 are related with other bacterial PGMs, stem from an ancient gene duplication event and are both monophyletic. (**A**) Phylogenetic analysis of chlamydial CT295 and CT815 homologs in eggNOG COG1109 (annotated as “phosphomannomutase”; 456 aligned positions; http://eggnogapi45.embl.de/nog_data/text/fasta/COG1109) clustered at 80% amino acid identity over 90% query coverage (n=2,829). Clades are colored by their affiliation to bacteria, eukaryota, and archaea in green, violet, and blue, respectively. Orange indicates the chlamydial CT295 clade (bacterial eggNOG ENOG4107QSU), and yellow indicates the chlamydial CT815 clade (bacterial eggNOG ENOG4107QJF). Maximum likelihood phylogenetic trees with best fit model LG+C60+G+F with 1,000 ultrafast bootstraps are shown. Bootstrap support for monophyly of chlamydial clades in the tree is ≥ 95%, and the SH-like approximate likelihood ratio is ≥ 80%. (**B**) Phylogenetic analysis of chlamydial CT295 homologs (bacterial eggNOG ENOG4107QSU; n=104) with its sister clade from COG1109 (575 aligned positions) used as outgroup (n=93). Maximum likelihood phylogenetic tree based on LG+C60+G+F best tree using the posterior mean site frequency model and 100 non-parametric bootstraps. Bootstraps ≥ 70% are indicated by filled circles at splits in the tree. Scale bar indicates 0.2 substitution per position.

**Figure 2: Figure supplement 2.**
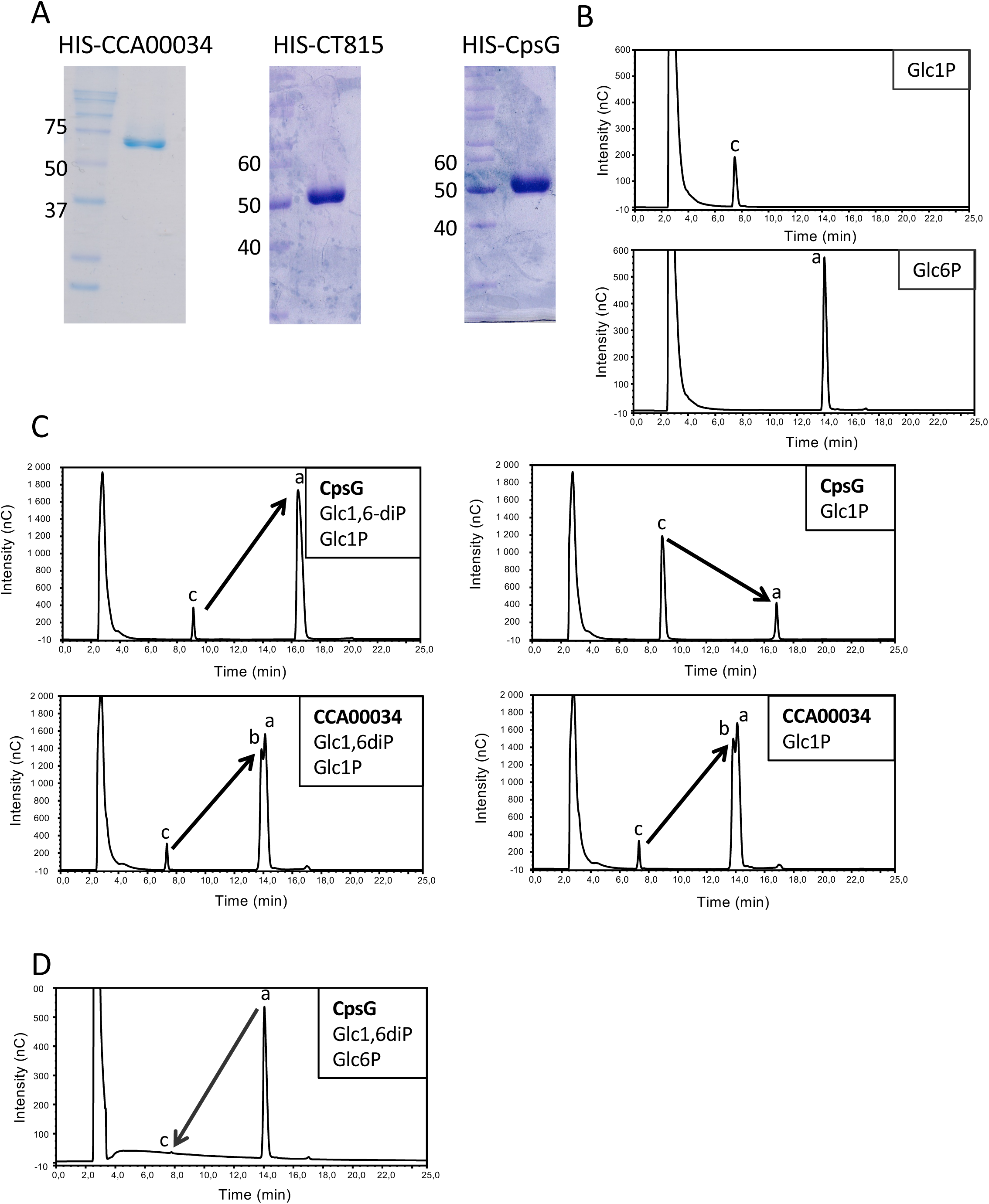
A – Migration profile of purified proteins. Purified His-tagged proteins were run on SDS-PAGE and stained with Coomassie blue. Molecular weights are indicated. B – Elution profiles of -Glc1P (peak c) and α−Glc6P (peak a). C – HIS-CpsG (top) or HIS-CCA00034 (bottom) were incubated with Glc1P in the presence (left) or absence (right) of the co-factor Glc1,6diP. The substrates and reaction products were identified by HPAEC-PAD. CpsG required the presence of Glc1,6diP for optimal conversion of Glc1P into Glc6P, while CCA00034 did not. Also, CCA00034 favored the conversion of of α-Glc6P (peak a) into β-Glc6P (peak b), while CpsG did not. D-HIS-CpsG was incubated with Glc6P in the presence of Glc1,6diP. Conversion of Glc6P into Glc1P was hardly detectable.

